# Peripheral *Taenia* infection increases immunoglobulins in the central nervous system

**DOI:** 10.1101/2020.09.26.314567

**Authors:** Sylvia Van Belle, Anja de Lange, Hayley Tomes, Rodney Lucas, Vinogran Naidoo, Joseph Valentino Raimondo

**Affiliations:** Division of Cellular, Nutritional and Physiological Sciences, Department of Human Biology, Neuroscience Institute and Institute of Infectious Disease and Molecular Medicine, Faculty of Health Sciences, University of Cape Town, Cape Town, South Africa

**Author notes:** Correspondence: Joseph V. Raimondo.

**Keywords:** Helminth, *Taenia crassiceps*, cysticercosis, immunoglobulin, blood-brain barrier, seizures

## Abstract

Human cysticercosis is a disease caused by larvae of the cestode *Taenia solium*. It is the most common cause of adult-acquired epilepsy world-wide where it exacts a debilitating toll on the health and well-being of affected communities. It is commonly assumed that the major symptoms associated with cysticercosis are a result of the direct presence of larvae in the brain. As a result, the possible effect of peripherally located larvae on the central nervous system are not well understood. To address this question, we utilised the *Taenia crassiceps* intra-peritoneal murine model of cysticercosis, where larvae are restricted to the peritoneal cavity. In this model, previous research has observed behavioural changes in rodents but not the development of seizures. Here we used ELISAs, immunoblotting and the Evans Blue test for blood-brain barrier permeability to explore the central effects of peripheral infection of mice with *Taenia crassiceps*. We identified high levels of parasite-targeting immunoglobulins in the serum of *Taenia crassiceps* infected mice. We show that the *Taenia crassciceps* larvae themselves also contain and release host immunoglobulins over time. Additionally, we describe, for the first time, significantly increased levels of IgG within the hippocampi of infected mice, which are accompanied by changes in blood-brain barrier permeability. However, these *Taenia crassiceps* induced changes were not accompanied by alterations to the levels of proinflammatory, pro-seizure cytokines in the hippocampus. These findings contribute to the understanding of systemic and neuroimmune responses in the *Taenia crassiceps* model of cysticercosis, with implications for the pathogenesis of human cysticercosis.

Graphical abstract created with BioRender.com

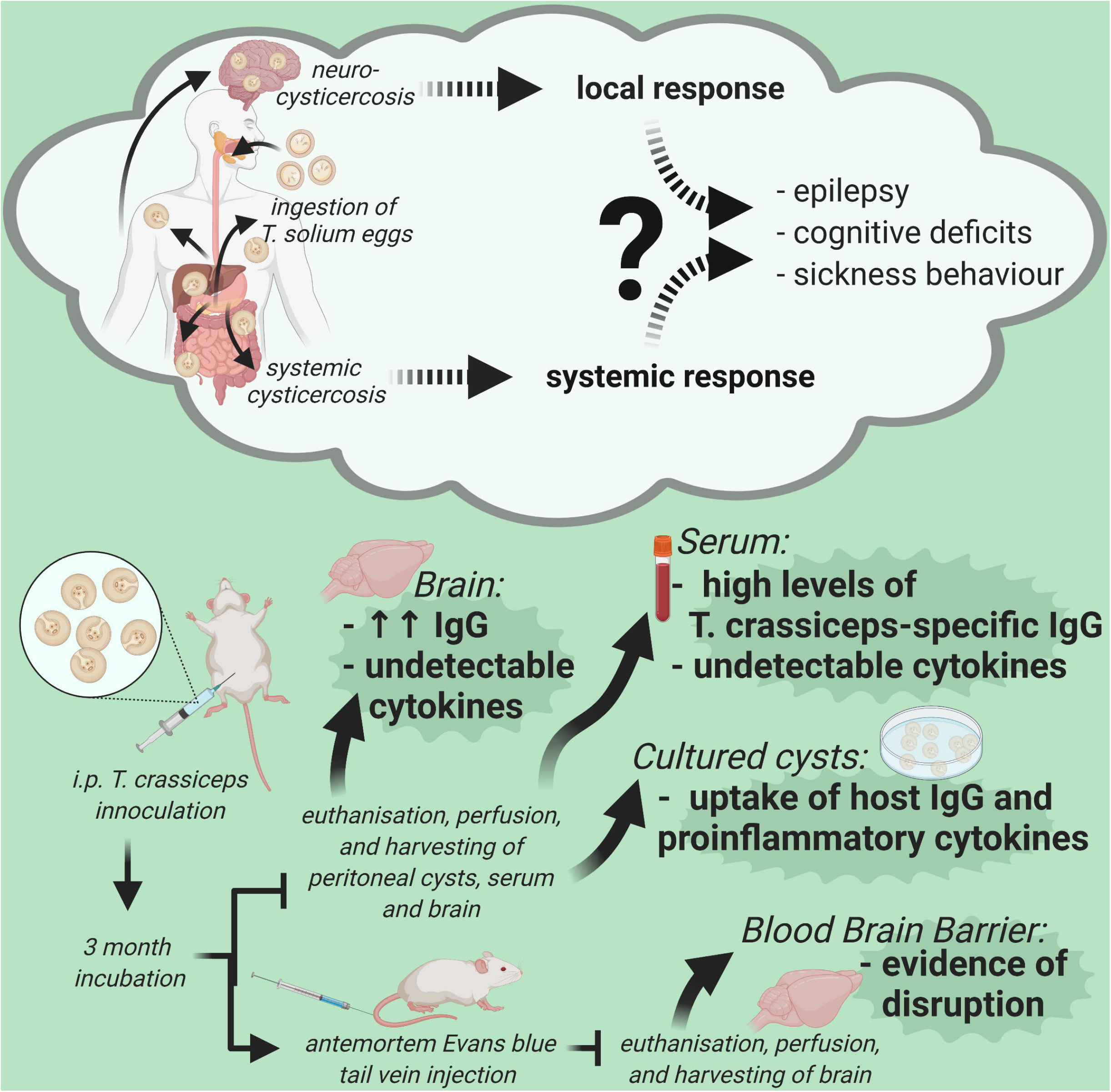

## 1 Introduction

Human cysticercosis is a common debilitating disease caused by larvae of the cestode *Taenia solium* (*T. solium*). An outdated conservative estimate suggests that up to 50 million people worldwide have cysticercosis (CDC, 1993). It is particularly common in rural or peri-urban areas of central and South America, Sub-Saharan Africa, India and Asia (White, 2018). Humans are the definitive host of the adult lifestage of the cestode *T. solium*. Cysticercosis occurs when humans act as accidental intermediate hosts (instead of pigs) following ingestion of the eggs in faeces from an infected host. The eggs change into stage one larvae after exposure to gastric acid, translocate across the intestinal epithelium and travel to in various tissues, with a strong predilection for the central nervous system (CNS). This results in what is termed neurocysticercosis. The major symptom associated with cysticercosis in humans is recurrent seizures. Cysticercosis is the most common cause of adult-acquired epilepsy world-wide; in *T. solium* endemic countries, approximately 29% of patients with epilepsy have cysticercosis (Ndimubanzi et al., 2010). In addition to seizures, patients with neurocysticercosis may also experience deficits in learning and memory (Nau et al., 2018). Whether this is a result of a direct function of the parasite, an immune response, recurrent seizures, or a combination of these, is uncertain.

It is generally assumed that the symptoms associated with neurocysticercosis are a result of the direct presence of larvae in the brain and the resulting pericystic inflammatory host response (Cangalaya et al., 2016). There are, however, interesting precedents for generalised parasitic infections having effects on the brain. For example, nodding syndrome caused by the parasite *Onchocerca volvulus*, which infects the skin and connective tissue without entering the brain, is characterised by head dropping and other seizure-like activity together with learning deficits (Lagoro and Arony, 2017). Interestingly the parasite does not invade the nervous system, and symptoms are due to host-generated antibodies against the parasite having a central, autoimmune, neurotoxic effect (Johnson et al., 2017). Furthermore, there is evidence that infection by other helminths, such as *Ascaris lumbricoides, Trichuris trichiura* and *Schistosoma* subspecies, can affect learning and cognition in infected individuals without infection of the CNS (Ezeamama et al., 2018; Pabalan et al., 2018). In the study of cysticercosis, little consideration has been given to the possibility that a systemic immune response to *Taenia* larvae, whether centrally or peripherally located, could be responsible for, or contribute to, neurological symptoms.

A valuable model system in which to explore this possibility is one in which mice are infected intraperitoneally with larvae of the cestode *T. crassiceps. T. crassiceps* is a related and antigenically similar cestode to *T. solium. T. crassiceps* typically infect canids, mustelids and felids as definitive hosts and various rodents as intermediate hosts (Willms and Zurabian, 2010). However, human infection as an accidental intermediate host has also been observed (Ntoukas et al., 2013). In the intra-peritoneal murine *T. crassiceps* model, larvae remain within the peritoneal cavity and do not invade the nervous system. As a result, it is a useful model system for studying potential effects of systemic responses to *Taenia* larvae on the brain (de Lange et al., 2019).

Previous research using this model has shown that, following infection, *Taenia* larvae are able to shift an initial, protective, systemic T helper type 1 immune response toward a T-helper type 2 response, which is more permissive for chronic infection. This Th2-type response is associated with high serum levels of cytokines IL-4, IL-5, IL-10 and antibodies IgG1, IgG4 and IgE (Dissanayake et al., 2004; Terrazas et al., 2010). To modulate the host immune response, *Taenia* larvae produce and excrete/secrete molecules which impair dendritic cell maturation and promote Th2-driving ability (Terrazas et al., 2010). Furthermore, the larvae can sequester and dispose of host immune proteins, including IgG (Flores-Bautista et al., 2018). The effect of these systemic parasite-induced host immune responses on the brain is still relatively uncertain. However, one study utilised the murine intra-peritoneal *T. crassiceps* infection model and reported behavioural changes, including impaired learning and memory, without larval invasion of the nervous system (Morales-Montor et al., 2014). This was found to be associated with variable changes in cytokine mRNA, for example relatively elevated mRNA levels of TNF-α and IL-6, but no change in IL-1β mRNA expression in the hippocampi of infected mice. One mechanism through which peripheral larvae may affect the brain could be via host-generated immunoglobulins against the larvae entering the nervous system via disruptions in the blood-brain barrier.

In this study we set out to investigate possible mechanisms by which a peripheral infection with *T. crassiceps*, and the resulting systemic immune response, could result in neurological changes such as those described by Montor et al. (2014). We employ the murine intra-peritoneal *T. crassiceps* infection model to elicit a systemic immune response, as demonstrated by high levels of parasite-targeting immunoglobulins in the serum of *T. crassiceps* infected mice. We describe, for the first time, significantly higher levels of IgG within the hippocampi of infected mice which are accompanied by changes in blood-brain barrier permeability. Unexpectedly, these *T. crassiceps-induced* alterations were not accompanied by changes in the levels of proinflammatory cytokines in the hippocampus. We further show that the *T. crassciceps* larvae contain host immunoglobulins and cytokines, and describe how these are released *in* vitro by the larvae over 3-10 days. These findings demonstrate that systemic host immune responses to infection with *Taenia* larvae can result in alterations in the brain, and thereby contribute to a wider understanding of the neuropathology in cysticercosis.

## 2 Materials and Methods

### 2.1 Animals

#### 2.1.1 Ethics

Mice were closely monitored for signs of distress (at least once a day), and all animal handling, care and procedures were carried out in accordance with South African national guidelines (South African National Standard: The care and use of animals for scientific purposes, 2008) and with authorisation from the University of Cape Town Animal Ethics Committee (Protocol No: AEC 015/015, AEC 019/025).

#### 2.1.2 Mice

In accordance with experiments conducted by Mahanty et al. (2013), female C57BL/6 mice were housed in the animal care unit at the Faculty of Health Sciences, University of Cape Town, under controlled temperature conditions and a 12-hour dark-light cycle. Control mice were housed in separate cages in the same facility.

#### 2.1.3 Infection

In accordance with the model described by Morales-Montor et al. (2014), 5-8 week old female C57BL/6 mice each received a single intra-peritoneal injection of 20 non-budding *T. crassiceps* larvae (ORF strain). Mice were killed 12-14 weeks later and the larvae harvested from the peritoneal cavity. Each mouse harboured between 200-500 larvae. Age-matched control mice were injected with phosphate-buffered saline (PBS) (**Figure 1**).

**Figure 1:**
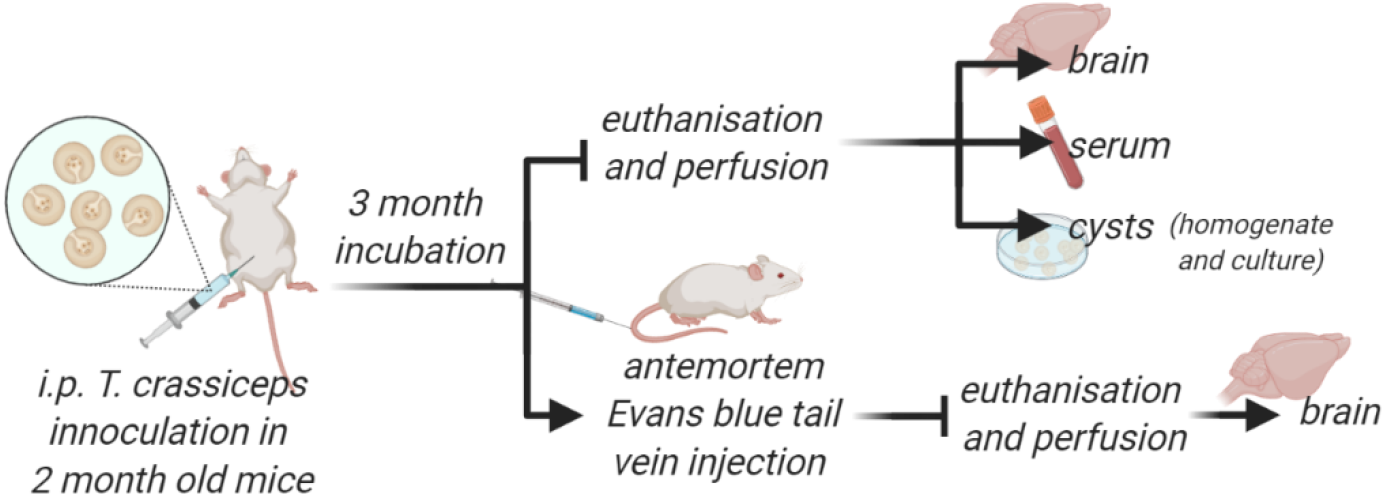
outline of infection timeline and sample collection protocol.

#### 2.1.4 Euthanisation

Mice were euthanised by halothane overdose followed by cervical dislocation.

### 2.2 Sample preparation

#### 2.2.1 Preparation of *Taenia crassiceps* whole cyst homogenate

Larvae were frozen at −80 °C after harvesting and washing. Upon thawing, larvae were suspended in a volume of PBS (1X, pH 7.4) threefold that of the larval volume. A protease inhibitor cocktail was added to this suspension (1% vol/vol, Sigma-Aldrich). The larvae were then homogenised on ice using a glass tissue homogeniser. The resulting mixture was centrifuged at 1 500 x g for 20 min at 4 °C. The supernatant, excluding the low density white floating layer, was collected, sterile filtered through a 0.22 μm size filter (Millex-GV syringe filter, Merck), and aliquoted and stored at −80 °C until use.

#### 2.2.2 Preparation of *Taenia crassiceps* excretory/secretory products

After harvesting, approximately 15 ml of PBS-washed larvae (+/- 15ml) was transferred into a culture flask with 50 ml culture medium (Earle’s Balanced Salt Solution with glucose (5.3 g/liter), Glutamax (1X), penicillin (50 U/ml), streptomycin (50 μg/ml), gentamicin sulphate (100 μg/ml) and nystatin (11.4 U/ml)). Larvae were maintained at 37 °C in 5 % CO_2_.

##### 2.2.2.1 Pooled excretory/secretory product collection

After 24 hours the medium was discarded and replaced with 50 ml fresh medium. Medium was collected every 3-5 days (and temporarily stored at −20 °C) and replaced with fresh medium for up to 20 days. After 20 days all the collected media was pooled and concentrated/buffer exchanged to PBS (1X, pH 7.4) using an Amicon stirred cell (Merck) with a 3 kDa molecular weight cut-off membrane.

##### 2.2.2.2 Excretory/secretory product collection separated by day

Medium was collected and replaced with 50 ml fresh medium at the end of day 1 *in vitro*, and again at the end of day 3, day 5, day 8 and day 10 in vitro. Each medium collection was concentrated/buffer exchanged to PBS (1X, pH 7.4) using an Amicon stirred cell (Merck) with a 3 kDa molecular weight cut-off membrane. This protocol was performed on two separate harvests of *T. crassiceps* larvae.

#### 2.2.3 Serum

Blood was withdrawn prior to transcardial perfusion and allowed to clot undisturbed at room temperature for 20 min. Samples were centrifuged at 1 500 x g for 10 minutes at 4°C. The supernatant was removed and stored at −80°C until use. Sera from infected (*n* = 5) and control mice (*n* = 5) were aggregated to provide adequate volumes for ELISAs and immunoblots.

#### 2.2.4 Hippocampal tissue

Mice were perfused with ice-cold PBS and both hippocampi were collected from control (*n* = 6) and infected (*n* = 8) mice. These samples were then frozen in liquid nitrogen and stored at −80 °C. Each hippocampus was thawed in 400 μl RIPA buffer with 0.2 % HALT™ Protease Inhibitor Cocktail (ThermoFisher Scientific) and sonicated for 15 s on ice. Samples were then centrifuged at 17 200 x g for 30 min at 4 °C and supernatants were saved and stored at −80 °C.

#### 2.2.5 Evan’s Blue injections

A 0.5 % solution of Evans Blue dye in PBS was prepared and filtered through Sartorius™ grade 3-HW smooth filter paper discs (ThermoFisher Scientific). Thirty minutes before euthanasia, each mouse received 200 μl of Evans Blue solution, injected into the lateral tail vein of infected (*n* = 5) and control (*n* = 3) mice. Animals were overdosed with halothane and transcardially perfused with ice-cold PBS. The hippocampi, frontal and remaining cortices and cerebellum were collected, dried and weighed. 1:3 w/v 50 % trichloroacetic acid (TCA)-0.9 % saline was added to each sample, and they were sonicated for 15s on ice. Samples were centrifuged at 10 000 x g for 20 min at 4 °C, and the supernatant was collected and diluted 1:3 in ethanol, before being stored at −80 °C (Kaya and Ahishali, 2011; Radu and Chernoff, 2013).

### 2.3 Experimental Procedures

#### 2.3.1 ELISAs

Levels of the inflammatory cytokines IL-1β, IL-6 and TNF-α were measured in supernatants from brain homogenates, serum and *T. crassiceps* excretory/secretory (E/S) product using commercially available ELISA reagents according to the manufacturer’s instructions (R&D Systems, Germany). 96 well plates were coated with 50 μl antibody solution (IL-1β: 3 μg/ml; IL-6: 3 μg/ml; TNF-α: 1 μg/ml), blocked with 4 % BSA and incubated with 50 μl of each sample. They were then incubated with biotinylated detection antibodies (50 μl of 0.3 μl/ml antibody solution), before adding streptavidin alkaline phosphatase solution. The phosphatase substrate was added 45 min before absorbance measurement. Absorbances were measure at 405nm with a reference wavelength of 492nm using a Glomax Explorer Microplate Reader (Promega), and unknown sample concentrations were interpolated from a standard curve from samples consisting of serial dilutions of recombinant cytokines. A detailed version of the protocol followed is available online (<u>dx.doi.org/10.17504/protocols.io.bh2fj8bn</u>). Where samples fell below the detection limit of ELISAs, they were allocated the value of the lower detection limit for the purposes of statistical comparison.

#### 2.3.2 Western blotting

##### 2.3.2.1 General Protocol

Hippocampal protein concentrations were determined according to the Pierce™ BCA Protein Assay (Thermo Fisher Scientific). Aliquots (20 μg) of the protein samples were denatured in SDS at 100°C, separated by electrophoresis on 12% SDS-PAGE, and then blotted to nitrocellulose membranes. Protein transfer was confirmed with Ponceau S stain (0.1% w/v Sigma-Aldrich). Membranes were blocked for 5 minutes using undiluted EveryBlot Blocking Buffer (Bio-Rad) and then incubated with primary antibodies (specified below) overnight in 4°C. Membranes were then treated for 1 h at room temperature with horseradish peroxidase-conjugated secondary antibodies, and imaged using a SynGene G:Box.

##### 2.3.2.2 Serum Taenia crassiceps-specific IgG detection

Two 12% gels were each loaded with 15 μg pooled E/S product and 15 μg cyst homogenate and were transferred to individual nitrocellulose membranes. Each membrane was then incubated overnight at 4 °C in 500 μl serum from either *T. crassiceps*-infected mice or control mice, such that any *T. crassiceps-specific* antibodies in the serum would bind to the antigens separated by electrophoresis in the E/S product or homogenate samples. The membranes were then incubated in goat-anti-mouse IgG-HRP (Bio-Rad) and developed and digitally imaged for analysis.

##### 2.3.2.3 Excretory/secretory product IgG detection

E/S samples collected in two batches at days-in-culture 3, 5, 8 and 10 were transferred onto a nitrocellulose membrane, incubated in a rabbit anti-mouse-IgG IgG (Abcam) and rabbit anti-beta-actin IgG (Abcam) mixture, and then in donkey-anti-rabbit IgG-HRP (Abcam), before the membrane was developed and digitally imaged for analysis.

##### 2.3.2.4 Hippocampal and serum IgG detection

Hippocampal protein aliquots (20 μg) from infected and control animals were loaded onto SDS-polyacrylamide gels. A single sample from an infected mouse was repeated on every blot for the purpose of data normalisation. For comparison of systemic IgG levels, another gel was loaded with serial dilutions of serum in RIPA buffer (0.5 μg, 0.05 μg and 0.005 μg serum) from infected mice and control mice, alongside 20 μg of the same hippocampal sample used for blot normalisation. All membranes were then incubated with antibodies as described in **Section 2.3.2.3**, developed and digitally imaged for analysis.

#### 2.3.3 Evans Blue assessment of blood-brain barrier integrity

Samples ample were diluted in 95% EtOH and a standard curve contained 0.05-100 μg/ml Evans Blue dye in 50 % TCA and 95% EtOH. Fluorescence was measured at 620 nm excitation and 680 nm emission using a Glomax Explorer Microplate Reader (Promega).

### 2.4 Data analysis

For ELISAs, cytokine concentrations were calculated from absorbance readings using the Glomax Explorer Microplate Reader.

For IgG quantification, the optical densities for actin (43 kDa) and IgG (50 kDa) were determined for each exposure (30-180 s, in 30s intervals) using ImageJ. The IgG value was divided by the actin value, calculated as a ratio to the reference sample that remained the same across blots. The IgG amount in that reference sample was interpolated as an average across exposures from a concentration curve calculated in GraphPad Prism 6 from a blot of known concentrations (IgG from Abcam ab133470, lot concentration at 0.6 mg/ml). The IgG amount was then calculated for each value at each exposure. Colourisation and background removal for image creation was performed post-analysis.

Due to skewed data distributions, non-parametric Mann-Whitney U tests were performed to compare the infected versus the control mice in all experiments. A value of *P* < 0.05 was considered significant, and the Hodges-Lehmann difference between means was used in lieu of an effect size calculation. All graphs show the median with the interquartile range. GraphPad Prism v6.0 was used for data analysis and graph creation.

## 3 Results

### 3.1 Chronic peritoneal *Taenia crassiceps* infection induces a robust humeral immune response

All *T*. crassiceps mice in our study had typical larval burdens (200-500 larvae), confirming that our mice were susceptible to peritoneal infection by the parasite. The infection was confined to the peritoneal cavity as no larvae were present in any of the brain samples harvested. We first wished to confirm that there was a systemic immune response to the intra-peritoneal infection of mice with *T. crassiceps*. To do so, we used western blotting to compare the amount of IgG in the serum of infected and uninfected mice. IgG was markedly increased in the serum of infected mice (pooled from 5 animals) compared to control animals (pooled from 5 animals) **(Figure 2a)**.

**Figure 2:**
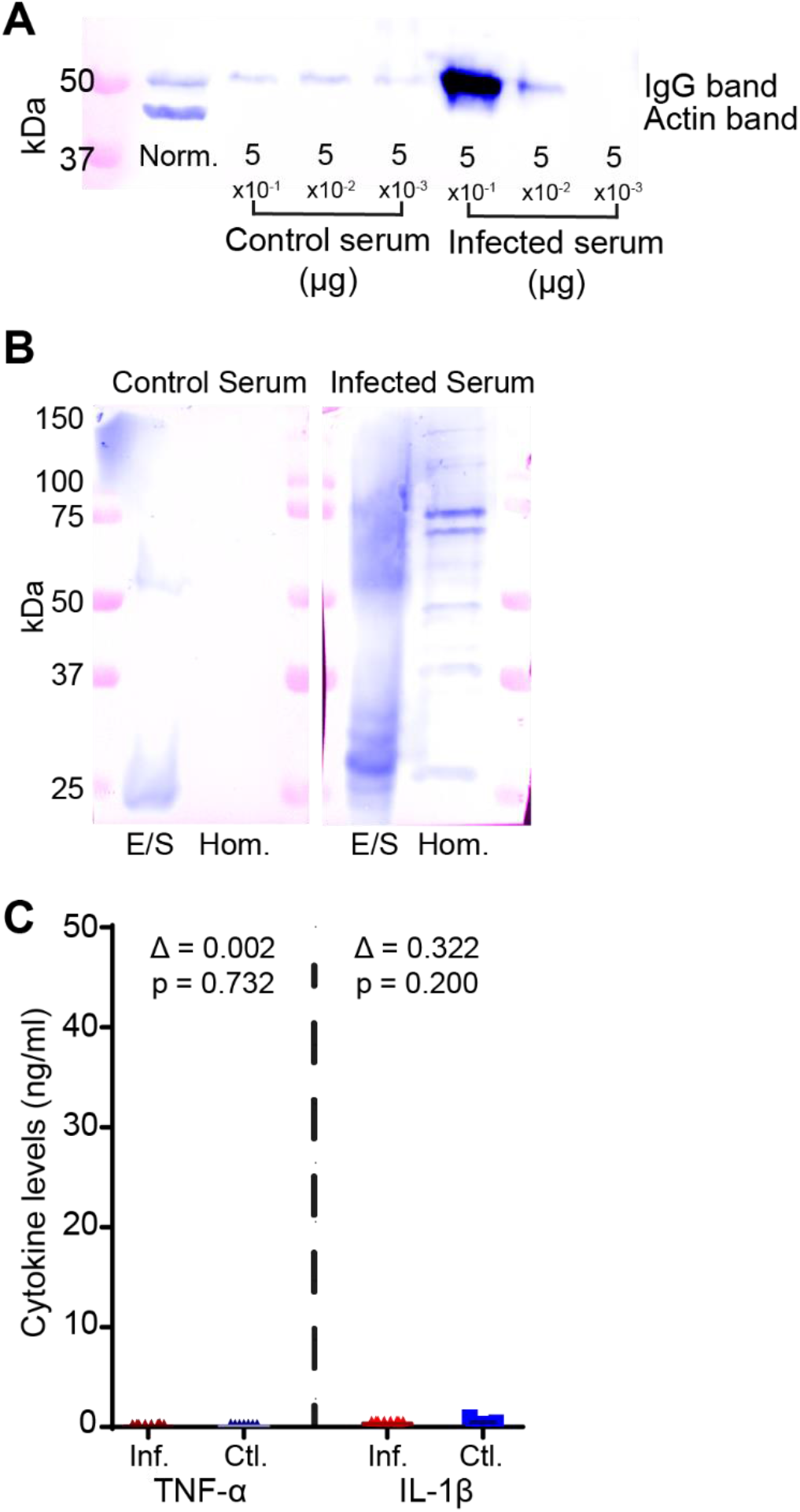
Chronic peritoneal *T. crassiceps* infection induces a robust humeral immune response. (**a**) IgG levels in serial dilutions of control versus infected serum (amount loaded in μg shown below the bands) show high levels of IgG in the serum of infected mice. (**b**) Two membranes each containing a lane run with *T. crassiceps* pooled E/S product and one run with larval homogenate. These were incubated in serum from infected mice (right) and serum from control mice (left) and probed with anti-mouse IgG. This demonstrates the presence of *T. crassiceps* specific antibodies in the serum of infected animals. (**c**) There was no significant difference in the serum cytokine levels of TNF α and IL-1β between infected and control animals. In addition, most serum samples had below-detectable levels of these cytokines (0.0078 pg/ml for TNF α and 0.0313 pg/ml for IL-1β). Norm. = normalisation sample (20 μg hippocampal homogenate from a *T. crassiceps* infected mouse); Inf. = Infected; Ctl. = control; E/S = Excretory/Secretory products; Hom. = larval homogenate. Δ = Hodges-Lehmann difference between the medians.

Next, we sought to determine whether infected mice might have raised specific antibodies circulating in their serum against antigens in the *T. crassiceps* antigens from the larval homogenate or pooled E/S product. To do so, two polyacrylamide gels were each loaded with *T. crassiceps* homogenate and E/S product and transferred onto nitrocellulose membranes. One membrane was then incubated with infected serum, while the other was incubated in serum from a control mice. Both were then probed with anti-mouse-IgG primary antibodies. These blots clearly show specific binding of serum IgG to multiple different antigens of both larval homogenate and E/S product in *T. crassiceps-infected* mice (**Figure 2b**).

We next sought to determine the serum levels of the proinflammatory cytokines TNF α and IL-1β which have known roles in epileptogenesis (Vezzani et al., 2016), using enzyme-linked immunosorbent assays (ELISAs). We did not detect significant differences in serum cytokine levels between infected and control mice **(Figure 2c**) with the values from the majority of the samples were below the detection threshold of 0.0078 pg/ml for TNF α (6/8 infected mice and 5/7 control mice were below 0.0078 pg/ml TNF α) and 0.0313 pg/ml for IL-1β (8/8 infected samples and 5/7 control mice were below 0.313 pg/ml IL-1β).

### 3.2 *Taenia crassiceps* larvae release host IgG and inflammatory cytokines over time

Given previous reports that *T. crassiceps* larvae can sequester and release host IgG (Flores-Bautista et al., 2018), we next sought to corroborate this observation and to further determine whether the larvae release host cytokines. To do so we evaluated the larval E/S product for IgG and cytokines TNF α, IL-1β and IL6 following 3, 5, 8 and 10 days in culture (see **Figure 3**). Western blots probing the *T. crassiceps* larval homogenate ande E/S product for mouse IgG showed clear evidence of host IgG, which steadily decreased over time **(Figure 3a,b**). After 10 days in culture, mouse IgG could nevertheless still be detected in the larval E/S product. Moreover, although there was variation in the total cytokine levels between the two batches of *T. crassiceps* larvae cultures, we were able to reliably detect the presence of the cytokines TNF α, IL-1β and IL-6 in the E/S product at all time points tested using ELISAs (**Figure 3c**).

**Figure 3:**
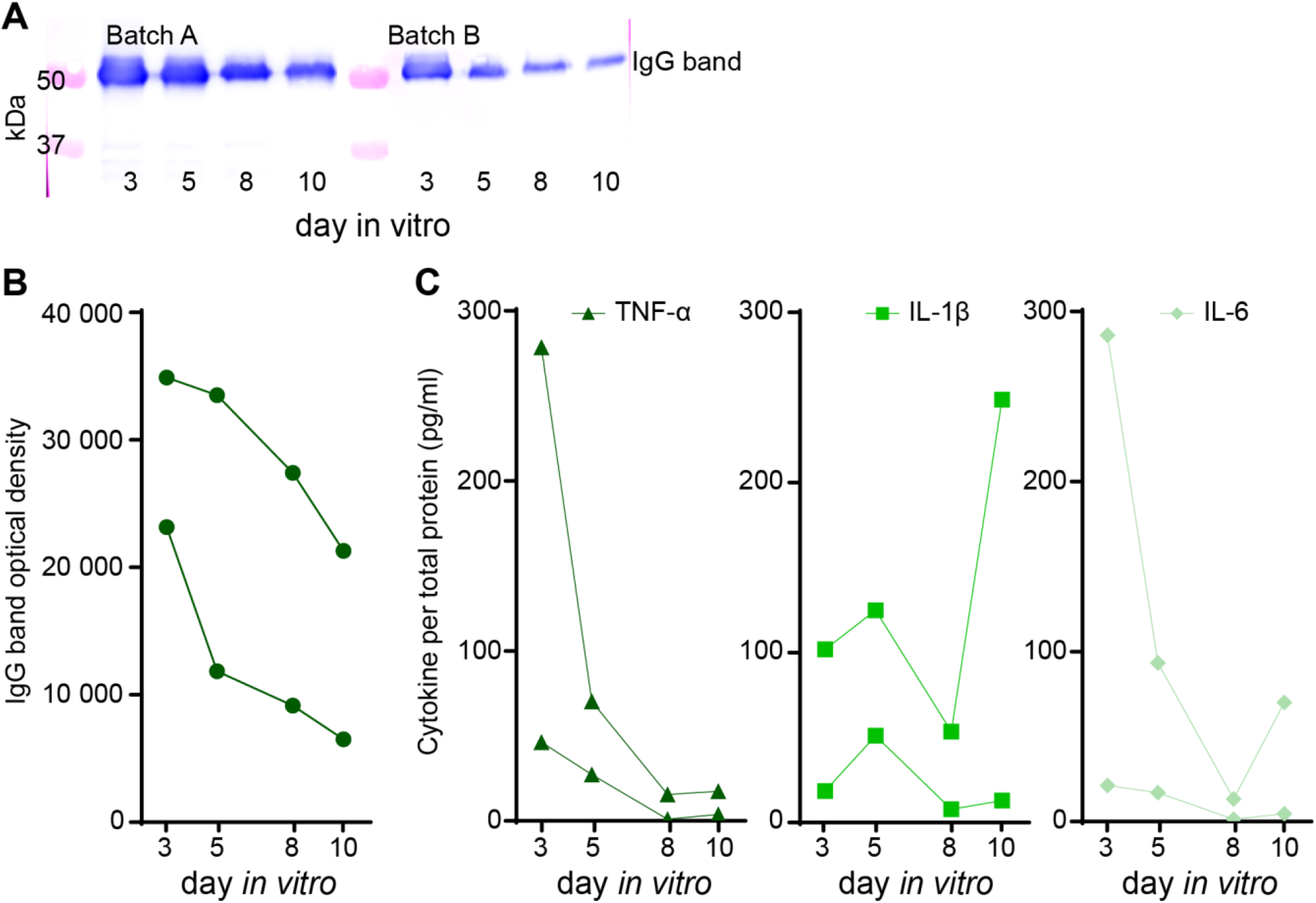
*T. crassiceps* larvae release host proteins over time. Culture medium from two separate batches of *T. crassiceps* larvae were collected on days 3, 5, 8 and 10 in culture. (**a**) Host IgG was detected in the larval E/S product, with levels decreasing with extended time in culture. (**b**) Measured optical density from blots shown in ‘a’, no proteins usable as a loading control were present for quantification. (**c**) The cytokines TNF-α, IL-1β and IL-6 could be detected in the E/S product at all time points tested.

### 3.3 Chronic peritoneal *Taenia crassiceps* infection increases intra-hippocampal IgG levels

Noting the clear systemic antibody response in **Figure 2**, we next wanted to determine if mice with intra-peritoneal *T. crassiceps* infection had higher levels of IgG within the brain parenchyma itself. All mice were PBS-perfused prior to dissection to prevent interference from intravascular IgG within the hippocampi and hippocampal samples from infected and control mice were immunoblotted for IgG. There was a striking difference between infected (*n* = 16) and control (*n* = 12) samples, with no IgG detectable in any of the control samples, and a clear presence of IgG present in most of the samples from infected mice (**Figure 4a**). Quantification of hippocampal IgG was performed using a normalisation sample and compared to a separately measured calibration curve with known amounts of IgG. There was a highly significant difference in hippocampal IgG levels between infected and control mice ((infected mice had a median of 0.0198 (IQR 0.0049 – 0.0266) ng per 20 μg while control mice had a median of 0.0002 (IQR 0.0 – 0.0004) ng per 20 μg for control mice, *P* < 0.0001, Mann-Whitney test).).

**Figure 4:**
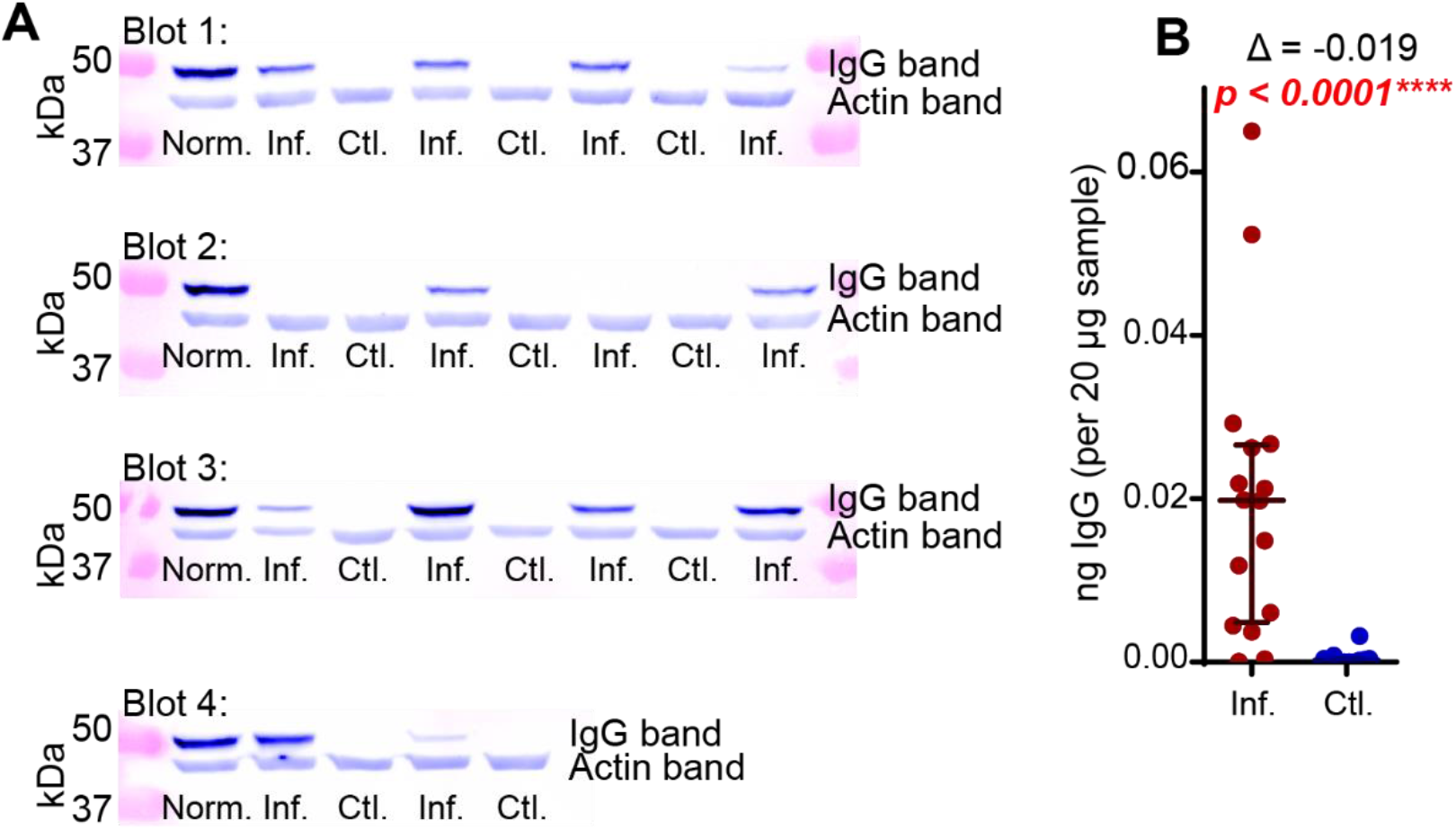
Chronic peritoneal *T. crassiceps* infection increases intra-hippocampal IgG levels. **a**. Western blots probing for IgG in homogenised hippocampi of infected (*n* = 16) and control (*n* = 12) mice (exposed for 3 minutes) (multiple blots needed to include all the samples). **b**. Population data of quantified hippocampal IgG levels from the blots in ‘**a**’. Inf.: Infected; Ctl.: Control; Norm.: infected hippocampal normalisation sample repeated between bots; Δ: Hodges-Lehmann difference between the medians.

### 3.4 Chronic peripheral *Taenia crassiceps* infection disrupts the blood-brain barrier but does not increase hippocampal proinflammatory cytokines

We next sought to determine a potential cause for the elevated intra-hippocampal IgG levels observed in infected mice (**Figure 4**), by injecting Evans Blue (EB) dye into the tail vein and tracked its presence within the brain parenchyma with fluorometry. This would inform us if peritoneal infection with *T. crassiceps* was also associated with blood-brain barrier dysfunction. We saw a clear trend in the amount of dye present in the brain, suggesting enhanced blood-brain barrier permeability in infected animals compared to controls. This was particularly evident within the cortex, with a median of 0.849 μg EB (IQR 0.797 – 1.230 μg) in the frontal cortex of infected mice compared to uninfected mice, with a median of 0.0.521 μg EB (IQR 0.055 – 0.630 μg), a Hodges-Lehmann difference of 0.732 μg EB (p = 0.008, Mann Whitney U test), and in the remaining cortex of infected mice there was a median of 1.213 μg EB (IQR 0.678 – 2.267 μg), versus a median of 0.623 μg EB (IQR 0.143 – 0.784 μg) in the uninfected mice, a Hodges-Lehmann difference of 0.591 μg EB (p = 0.032, Mann Whitney U test) (**Figure 5b**).

**Figure 5:**
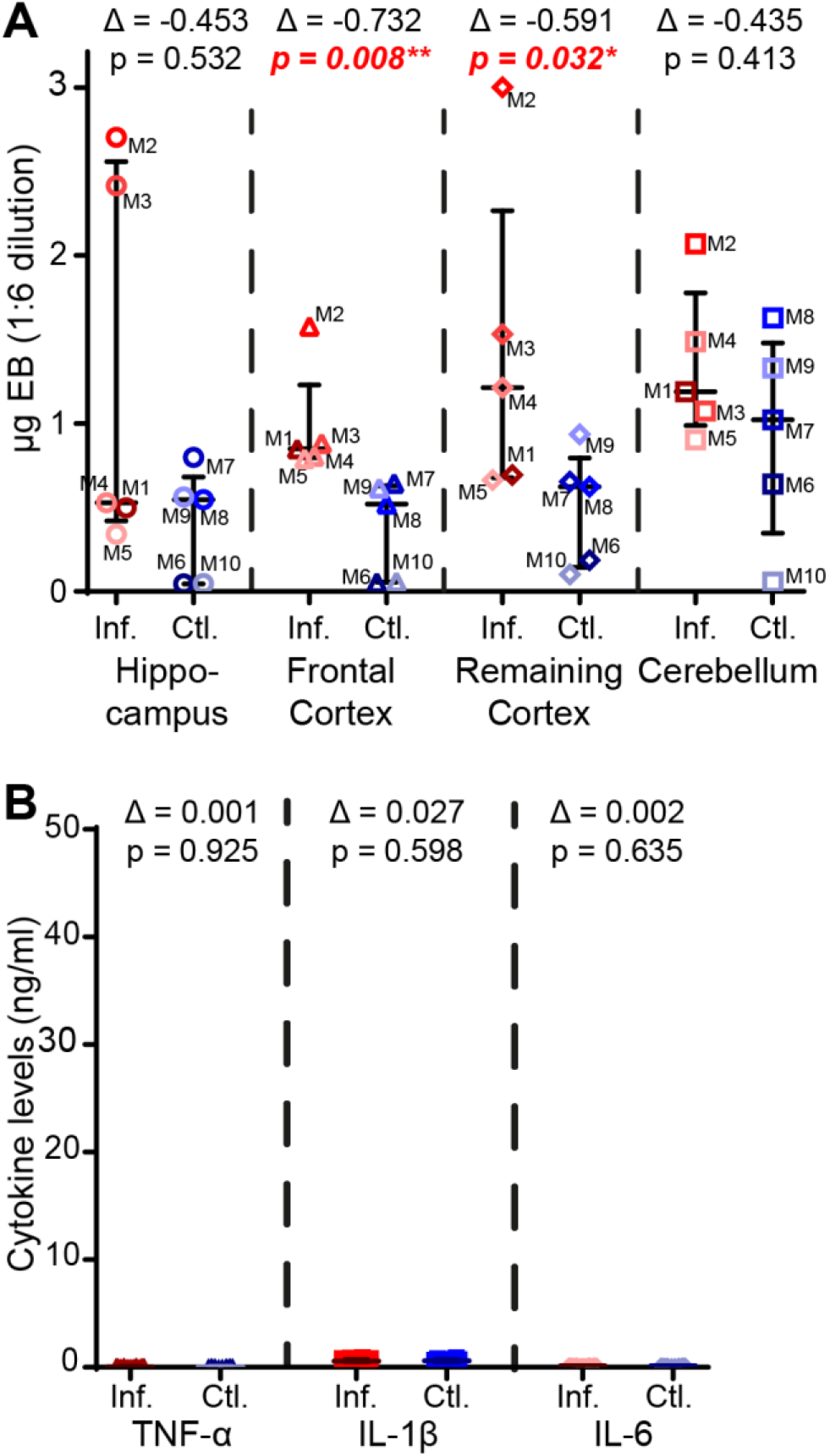
Chronic peripheral *T. crassiceps* infection disrupts the blood-brain barrier but does not increase hippocampal proinflammatory cytokines. **a**. Levels of Evans blue in homogenised samples from various areas in the brains of infected and uninfected mice. There was a trend towards an increased level of Evans Blue in the brains of infected mice, particularly in the cortex. **b**. Cytokine levels in the hippocampi were above the detection threshold, however, no difference between infected and control samples was observed. (Inf.: Infected; Ctl.: Control; EB: Evans Blue; Δ: Hodges-Lehmann difference between the medians.)

Finally, following our observation of an increase in intra-hippocampal IgG and a disruption in blood-brain barrier function in mice with peritoneal *T. crassiceps* infection, it is reasonable to assume that this could lead to enhanced parenchymal inflammation. This in turn could modulate network excitability. We therefore performed ELISAs of proinflammatory cytokines (TNF-α, IL-1β and IL-6), with previously described roles in neuroinflammation and epilepsy (Vezzani et al., 2016). Interestingly, the hippocampal cytokine levels were uniformly low and we did not detect significant differences in any of the three cytokines between infected and control mice (**Figure 5a**). The median hippocampal TNF-α levels (detection threshold of 0.0078 ng/ml) in infected compared to control mice were 0.0569 ng/ml (IQR 0.0341 – 0.0842 ng/ml) versus 0.0509 ng/ml (IQR 0.0404 – 0.0897 ng/ml) respectively, a Hodges-Lehmann difference of 0.0006 (p = 0.9250, Mann Whitney U test). The median hippocampal IL-1β levels (detection threshold of 0.0313 ng/ml) in infected versus control hippocampi were 0.5735 ng/ml (IQR 0.5058 – 0.7350 ng/ml) versus 0.6230 ng/ml (IQR 0.5185 – 0.7510 ng/ml) respectively, a Hodges-Lehmann difference of 0.0265 (p = 0.5984, Mann Whitney U test). Finally, the median IL-6 levels (detection threshold of 0.0039 ng/ml) were not significantly different in infected and control mice, with 0.1315 ng/ml (IQR 0.1128 – 1.388 ng/ml) versus 0.1320 ng/ml (0.1185–0.1400 ng/ml) respectively, a Hodges-Lehmann difference of 0.0020 (p = 0.6346, Mann Whitney U test).

## 4 Discussion

Here we used ELISAs, western blots and the Evans Blue test in a murine model of peripheral *T crassiceps* infection to investigate possible mechanisms by which a peripheral cysticercosis infection could result in neurological changes. We found high levels of parasite-targeting immunoglobulins in the serum of *T. crassiceps* infected mice. In addition, we found that the *T. crassiceps* larvae themselves also contain and release host immunoglobulins over time, indicating that a systemic immune response has been mounted. Furthermore, we observed significantly increased levels of IgG within the hippocampi of infected mice, which were accompanied by changes in blood-brain permeability. However, these *T. crassiceps* induced alterations were not accompanied by alterations to the levels of proinflammatory cytokines in the hippocampus, as described by previous authors (Morales-Montor et al., 2014), which could have contribute to behaviour changes and/or epileptogenesis.

We found that intra-peritoneal infection of C57BL/6 mice produced robust, and reliable chronic infections with approximately 500 cysts harvested from each mouse. This is in contrast to work suggesting that C57BL/6 mice are resistant to the ORF strain of *T. crassiceps* (Reyes et al., 2009). This likely points to significant variability in susceptibility to infection between strains or even perhaps sub-strains of mice. In our experiments, C57BL/6 mice were easily infected and mounted a significant humeral response to the parasite, as evidenced by the presence of *T. crassiceps* targeting antibodies in the serum. In addition, we were able to corroborate recent research, which has shown that the larvae themselves can take-up and release host protein, including IgG (Flores-Bautista et al., 2018). We found that even after 10 days in culture *T. crassiceps* larvae were still releasing host IgG and cytokines into the culture media. It is therefore important for those using *T. crassiceps* larvae derived homogenate or E/S products to investigate the modulation of host immune responses and to exercise caution as these products likely contain both host IgG and cytokines, which could influence their results.

Other research using the intra-peritoneal *T. crassiceps* infection model in Balb/c mice has shown that cysticercosis results in behavioural changes without larval invasion of the nervous system, including impairment in behavioural tasks such as the object recognition test and the forced swim test (Morales-Montor et al., 2014). It is well-known that peripheral infections in mammals are characterised by local, systemic and CNS effects. CNS effects may give rise to ‘sickness behaviour’ (Ghai et al., 2015) and IL-1β and TNF-α, produced particularly as part of the acute phase response, are thought to be important mediators of this neuro-immune signalling (Cartmell et al., 1999). In our experiments, following 12 weeks of infection, we found low to undetectable levels of IL-1β and TNF-α in serum, and no difference between control and infected animals. This is consistent with previous reports that although *T. crassiceps* infection results in a T helper type 1 proinflammatory immune response in the first 2 weeks following infection this is followed by a sustained T helper type 2 (Th2) response with low levels of IL-1β and TNF-α and high levels of IgG (Peon et al., 2013). This suggests that although IL-1β and TNF-α can cross the blood-brain barrier (Banks et al., 1995), centrally-acting serum IL-1β and TNF-α,are unlikely to be responsible for any sickness associated behaviours in the animals due to low serum levels following chronic *T. crassiceps* infection.

Interestingly, we found raised IgG in the hippocampi of infected mice, compared to control mice. It is possible that this IgG could have some functional effect through local immune activation (Kadota et al., 2000) although we did not demonstrate this here. In addition, our observation of raised parenchymal IgG was accompanied by changes in blood-brain barrier permeability. The blood-brain barrier, consisting of; negatively charged endothelial cells cross-linked through tight-junctions, smooth muscles cells and pericytes, ensures that only small molecules that are primarily lipophilic and cationic are able to cross from blood vessels into brain parenchyma without specialised transport channels (Finke and Banks, 2017; Ribeiro et al., 2012). This would normally exclude immunoglobulins from entering the brain tissue, however, blood-brain barrier breakdown has been shown to increase parenchymal IgG either via passive diffusion or via transmigration of immune cells such as plasma cells (Sweeney et al., 2019). It is uncertain whether the high levels of brain IgG observed in our study were a result of blood-brain barrier breakdown in infected animals, contributed to blood-brain barrier disruption, or both.

It is possible that the raised parenchymal IgG observed here could precipitate a local inflammatory response, including the local production of cytokines, with consequent effects on neuronal networks (Prieto and Cotman, 2017). However, we found low or undetectable levels of the cytokines IL-1β, TNF-α and IL-6, with no differences between infected and control animals. These findings contrast those of Lopez-Griego et al. (2015), who using quantitative PCR for cytokine mRNA found a modest but significant increase in TNF-α and IL-6 but not IL-1β mRNA in the hippocampi of infected female mice. This technique may be more sensitive than ELISAs, although to what degree mRNA expression relates to differences in translated cytokine levels is uncertain.

Given that IL-1β and TNF-α acting directly within the brain parenchyma have been strongly linked to changes in network excitability and seizures (Vezzani et al., 2011), our finding that there were no differences in these cytokines in the brains of *T. crassiceps* versus control animals is important in the context of epilepsy secondary to cysticercosis. Firstly, it may help explain why seizures are not observed in the *T. crassiceps* intra-peritoneal murine model of cysticercosis. Secondly, it supports the idea that people with cysticercosis, but with no larvae residing in the CNS, do not typically experience symptoms (Garcia et al., 2014). This suggests that for precipitation of seizures in cysticercosis, it is likely a necessary condition that the larvae must be directly located within the nervous tissue itself. As our findings suggest, it is still possible that peripheral infection, by driving potential additional disruption to the blood-brain barrier could exacerbate the epileptogenic process in neurocysticercosis (Guerra-Giraldez et al., 2013; Marchi et al., 2007). Taken together our findings contribute to the understanding of neuroimmune responses in the *T. crassiceps* model of cysticercosis, and more broadly, disease processes in human cysticercosis.

## Acknowledgements

The ORF strain of Taenia crassiceps was generously provided by Siddartha Mahanty (University of Melbourne). The research leading to these results has received funding from a Royal Society Newton Advanced Fellowship (NA140170) and a University of Cape Town Start-up Emerging Researcher Award to JVR and grant support from the Blue Brain Project, the National Research Foundation of South Africa, Wellcome Trust and the FLAIR Fellowship Programme (FLR\R1\190829): a partnership between the African Academy of Sciences and the Royal Society funded by the UK Government’s Global Challenges Research Fund. AdL received financial support from the National Research Foundation (110743), the Oppenheimer Memorial Trust (20787/02) and the University of Cape Town (Doctoral Research Scholarship). The funders had no role in study design, data collection and analysis, decision to publish, or preparation of the manuscript. We thank Ed Sturrock and Sylva Schwager (University of Cape Town) for generously allowing us to use their SynGene G:Box.

## Notes

### Competing Interest Statement

The authors have declared no competing interest.

